# *Wolbachia pipientis* occurs in *Aedes aegypti* populations in New Mexico and Florida, USA

**DOI:** 10.1101/455014

**Authors:** Aditi Kulkarni, Wanqin Yu, Jinjin Jiang, Concepcion Sanchez, Ajit K. Karna, Kalli J.L. Martinez, Kathryn A. Hanley, Michaela Buenemann, Immo A. Hansen, Rui-de Xue, Paul Ettestad, Sandra Melman, Dagne Duguma, Mustapha Debboun, Jiannong Xu

## Abstract

The mosquitoes *Aedes aegypti* (L.) and *Ae. albopictus* Skuse are the major vectors of dengue, Zika, yellow fever and chikungunya viruses worldwide. *Wolbachia*, an endosymbiotic bacterium present in many insects, is being utilized in novel vector control strategies to manipulate mosquito life history and vector competence to curb virus transmission. Earlier studies have found that *Wolbachia* is commonly detected in *Ae*. *albopictus* but rarely detected in *Ae. aegypti*. In this study, we used a two-step PCR assay to detect *Wolbachia* in wild-collected samples of *Ae. aegypti.* The PCR products were sequenced to validate amplicons and identify *Wolbachia* strains. A loop-mediated isothermal amplification (LAMP) assay was developed and used for detecting *Wolbachia* in selected mosquito specimens as well. We found *Wolbachia* in 85/148 (57.4%) wild *Ae. aegypti* specimens from various cities in New Mexico and in 2/46 (4.3%) from St. Augustine, Florida. We did not detect *Wolbachia* in 94 samples of *Ae. aegypti* from Deer Park, Harris County, Texas. *Wolbachia* detected in *Ae. aegypti* from both New Mexico and Florida was the *w*AlbB strain of *Wolbachia pipientis.* A *Wolbachia* positive colony of *Ae. aegypti* was established from pupae collected in Las Cruces, New Mexico in 2018. The infected females of this strain transmitted *Wolbachia* to their progeny when crossed with males of Rockefeller strain of *Ae. aegypti*, which does not carry *Wolbachia.* In contrast, none of the progeny of progeny of Las Cruces males mated to Rockefeller females were infected with *Wolbachia*.

## 1 INTRODUCTION

*Wolbachia* are obligate intracellular bacteria found in a wide range of terrestrial arthropods and nematodes (Werren, Baldo, & Clark, 2008). The bacterium was discovered in the reproductive tissues (testes and ovaries) of the mosquito *Culex pipiens* L. by Hertig and Wolbach in 1924 (Hertig & Wolbach, 1924) and was formally described as *Wolbachia pipientis* by Hertig in 1936 (Hertig, 1936). About 60-70% of all insect species harbor *Wolbachia*, including some mosquito species (Hilgenboecker, Hammerstein, Schlattmann, Telschow, & Werren, 2008). *Wolbachia* can be a powerful reproductive manipulator, inducing cytoplasmic incompatibility (CI), parthenogenesis, feminization of males, and male killing in various host species (Werren et al., 2008). These properties have been exploited for development of *Wolbachia* as a novel strategy for vector mosquito control. *Wolbachia-*induced CI favors the reproductive success and spread of colonized females in populations, which can be used to drive desirable traits, including resistance to infection with vector-borne pathogens, into a population. On the other hand, infected mosquito males can cause CI in a population with the presence of different *Wolbachia* strains or no infection, which can be used for sterile insect technique (SIT) to decrease vector populations (Flores & O’Neill, 2018).

*Aedes (Stegomyia) aegypti* (L.) and *Ae. (Stegomyia) albopictus* Skuse are the major vectors for the transmission of several arthropod-borne viruses (arboviruses) among humans, particularly dengue, Zika, yellow fever, and chikungunya viruses. *Wolbachia* is commonly found in *Ae. albopictus* (de Albuquerque, Magalhaes, & Ayres, 2011; Joanne et al., 2015; Kitrayapong, Baimai, & O’Neill, 2002), but until recently *Ae. aegypti* was thought not to carry this bacterium (Gloria-Soria, Chiodo, & Powell, 2018; Kitrayapong et al., 2002; Kittayapong, Baisley, Baimai, & O’Neill, 2000). However, *Wolbachia* sequences were found in wild *Ae. aegypti* in a few recent investigations. *Wolbachia* 16S ribosomal RNA gene sequencing reads (operational taxonomic units, OTUs) were detected in *Ae. aegypti* larvae collected from Jacksonville, Florida (Coon, Brown, & Strand, 2016) as well as in two *Ae. aegypti* adult pools collected from Thailand (Thongsripong et al., 2017). More recently, *Wolbachia* 16S reads were detected in a few individuals of *Ae. aegypti* collected from Houston, Texas, though the regular PCR to amplify other *Wolbachia* genes was not successful (Hegde et al., 2018).

Establishing the prevalence of *Wolbachia* in *Ae. aegypti* is critical to public health, because over the past decade, *Ae. aegypti* transinfected with *Wolbachia* have been generated with the goal of blocking transmission of dengue virus (Bian, Xu, Lu, Xie, & Xi, 2010; Bull & Turelli, 2013; Hoffmann et al., 2011; McMeniman et al., 2009; O’Neill, 2018; Walker et al., 2011). This approach was initially aimed at shortening mosquito lifespan below the extrinsic incubation period of the virus (McMeniman et al., 2009), but in the course of these experiments it was discovered that transinfection of *Ae. aegypti* with *Wolbachia* strain *w*MelPop also blocks dengue and chikungunya virus infections of the mosquito (Moreira et al., 2009). A successful large field trial in Australia showed a stable establishment and slow but steady spread of released *Ae. aegypti* transinfected with *w*Mel in the study area (Schmidt et al., 2017). However, if a population of *Ae. aegypti* were to harbor an autochthonous strain of *Wolbachia*, then this native strain would have a high potential to prevent invasion of a virus-blocking strain that exhibits incompatibility with the native strain (Hoffmann, Ross, & Rasic, 2015). This effect was demonstrated in a study of *Ae. albopictus*, wherein the *w*Mel transinfected line produced complete bidirectional incompatibility with a wildtype line carrying *w*AlbA and *w*AlbB, with 0% hatch rate from crossing between females of either strain with males of the other strain (Blagrove, Arias-Goeta, Failloux, & Sinkins, 2012). On the other hand, complete CI could favor SIT for population reduction.

As part of a project to map the distribution of *Ae. aegypti* and *Ae. albopictus* in New Mexico in 2017 and characterize their mosquito microbiota, we unexpectedly detected *Wolbachia* 16S rRNA gene amplicon in wild-caught *Ae. aegypti* in Las Cruces, New Mexico. A more comprehensive survey was then conducted using a two-step PCR assay, which revealed a 57.4% prevalence of *Wolbachia* carriage in 148 specimens of *Ae. aegypti* collected from eight cities across New Mexico. *Wolbachia* was also detected in two of 48 specimens of *Ae. aegypti* from St. Augustine, Florida (4.2% prevalence), but not detected in 94 specimens of *Ae. aegypti* from Deer Park, Harris County, Texas. A *Wolbachia* infected *Ae. aegypti* strain was established from wild pupae collected from Las Cruces. The cross of the strain with the *Wolbachia* uninfected Rockefeller strain demonstrated maternal transmission of *Wolbachia* to progeny.

## 2 MATERIALS AND METHODS

### 2.1 Mosquito collections and species identification

*Aedes aegypti* and *Ae. albopictus* mosquitoes were collected using gravid and sentinel traps in New Mexico in 2017 by the SouthWest *Aedes* Research and Mapping (SWARM) project team, in Florida in 2016 by the Anastasia Mosquito Control District, and in Texas in 2018 by the Harris County Public Health Mosquito and Vector Control Division. The location details of the samples are presented in Table 1 and Figure 1. Although *Ae. aegypti* and *Ae. albopictus* were commonly collected in the same trap in Texas, they were never collected in the same trap in New Mexico or Florida. Mosquitoes were sorted and identified as *Ae. aegypti* or *Ae. albopictus* based on morphology. The species identity was confirmed by a species-diagnostic PCR assay that amplifies species-specific fragments of internal transcribed spacer 1 (ITS1) of ribosomal DNA, as described by (Higa, Toma, Tsuda, & Miyagi, 2010). The primers used in this study are listed in Table S1. The PCR was conducted using 2 × PCR master mix (MCLab, South San Francisco, CA) with ∼20 ng DNA, 0.2 µM primers, and cycling parameters as: 35 cycles of denaturing at 94°C for 15 seconds, annealing at 55°C for 15 seconds, and extension at 72°C for 30 seconds with extra 5 min in the last cycle for final extension.

**Figure 1.**
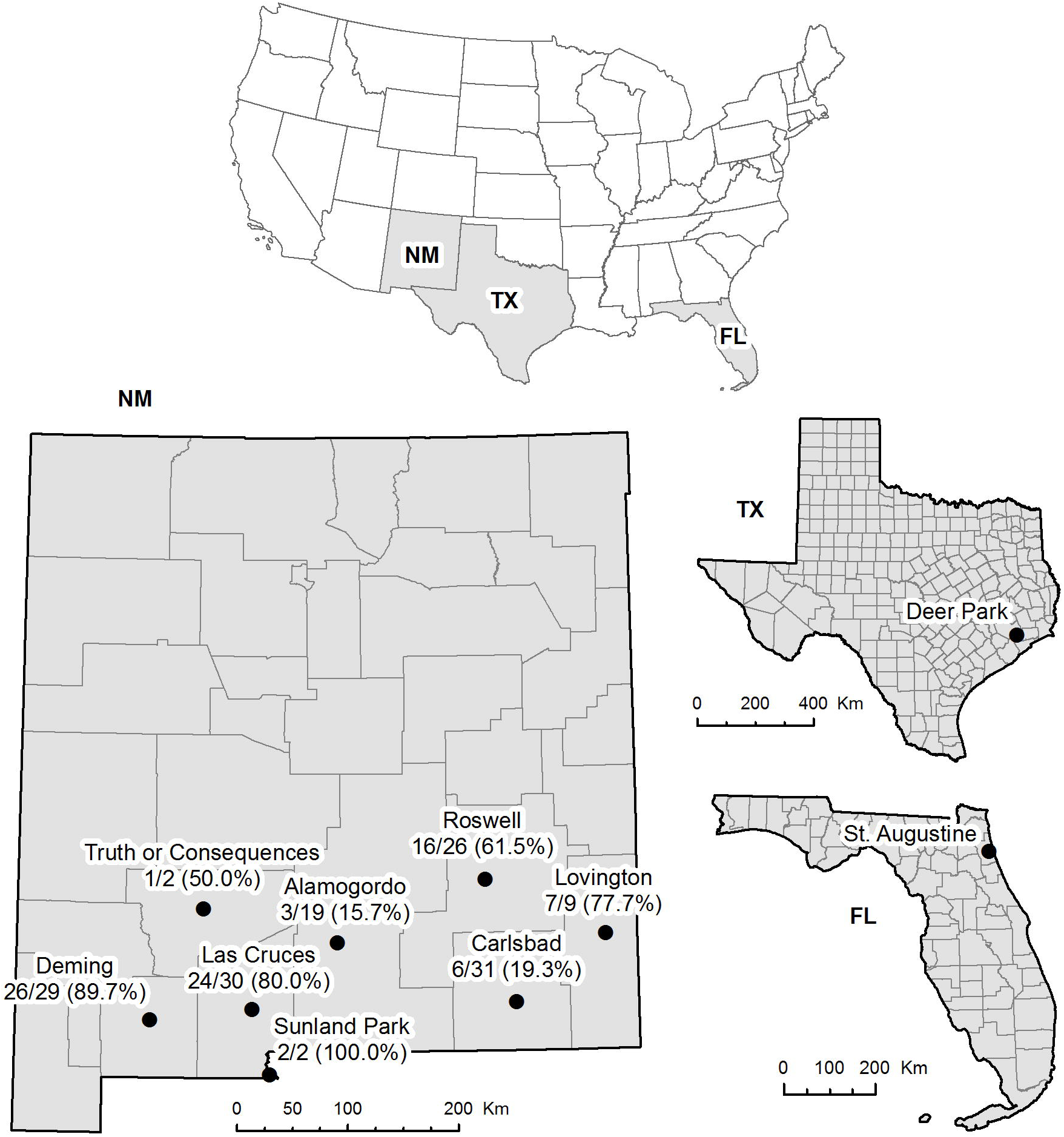
Maps of the sites where *Ae. aegypti* mosquitoes were sampled in this study. No. of *Wolbachia* positive/No. of tested (%) was displayed in sampling sites.

### 2.2 DNA isolation, Bacterial 16S rDNA PCR, cloning and sequencing

Mosquito specimens from traps were desiccated in most cases. For each mosquito specimen, the abdomen was separated from the thorax by pulling gently with tweezers that were cleaned with 75% ethanol between samples. Abdomens were used for detecting associated microbiota. Metagenomic DNA was isolated individually from each abdomen using DNAzol following the manufacturer’s protocol (Thermo Fisher Scientific, Waltham, MA). Briefly, one abdomen was homogenized in 100 µl DNAzol, and centrifuged at 12,000 ×g for 10 min. Supernatant was transferred to a new tube and 50 µl ethanol was added, and the tube was centrifuged at 12,000 ×g for 10 min for DNA precipitation. The DNA pellet was dissolved in 30 µl H2O. A bacterial 16S rDNA fragment covering V1 to V3 was amplified from individual DNA using primers 27F and 519R (Table S1), as we previously reported (Wang, Gilbreath, Kukutla, Yan, & Xu, 2011). PCR was run using 2× PCR master mix with 0.2 µM primers and cycling parameters: 35 cycles of denaturing at 94°C for 15 seconds, annealing at 52°C for 15 seconds, and extension at 72°C for 30 seconds with extra 5 min in the last cycle for final extension. PCR products were purified and cloned into the plasmid pMiniT 2.0 using a NEB PCR cloning kit (New England Biolabs, Ipswich, MA) following manufacturer’s instructions. Colony PCR was conducted to amplify the inserts using SP6 forward and T7 reverse primers. The PCR products with a size of ∼ 500 bp were sent for Sanger sequencing at a commercial provider (MCLab).

### 2.3 *Wolbachia* PCR assays and sequencing

The PCR assays using primer sets for the *Wolbachia gatB* and *ftsZ* gene from (Baldo et al., 2006) (Table S1), were used for detecting *Wolbachia* in mosquitoes. PCR was run using 2× PCR master mix (MCLab) with a primer concentration of 0.2 µM and the following cycling parameters: 35 cycles of denaturing at 94°C for 15 seconds, annealing at a temperature optimal for the amplicon (Table S1) for 15 seconds, and extension at 72°C for 30 seconds with an extra 5 min in the last cycle for final extension. For specimens that showed a faint band or no visible band after a *Wolbachia* target PCR, a second round PCR was conducted. The first round PCR product was diluted 100 times with H2O and 1 µl was used as template for the second PCR. In 85 *Wolbachia* positive specimens from NM, 8 showed a clear band in the first PCR and the remaining specimens showed a faint or no band in the first PCR but a clear band in the second PCR.

To validate the *Wolbachia* detection in *Ae. aegypti*, we designed primers to amplify fragments from two *Wolbachia* genes encoding phosphoesterase (PE) and diaminopimelate epimerase (DE) based on the draft genomes of *w*AlbB (Mavingui et al., 2012) and *w*AlbA (GenBank accession NWVK00000000.1). The sequences of the two genes have distinctive inter-strain differences, enabling the design of strain specific primers (Table S1). A subset of specimens that were positive from the first or second PCR were subjected to the validation PCR and sequencing. The products were sequenced at MCLab, and the sequences were deposited in GenBank; the accession numbers are presented in Table S2.

### 2.4 Loop-mediated isothermal amplification (LAMP) assay

Loop-mediated isothermal amplification (LAMP) was developed as an additional assay for the detection of *Wolbachia* in mosquitoes. Oligonucleotides for LAMP were designed using Primer Explorer V5 software available on the website (http://primerexplorer.jp/lampv5e/index.html). The sequences of oligos for the 16S rRNA gene are listed in Table S1. The LAMP reactions were conducted using a NEB LAMP kit with Bst 3.0 (M0374, NEB). The reaction mixture, consisting of 1X isothermal amplification buffer II, 6 mM MgSO4, 1.4 mM of each of the deoxynucleotide triphosphates (dNTPs), 1.6 µM Forward Inner Primer/Backward Inner Primer, 0.4 µM F3/B3 primers, 0.8 µM Loop Forward/Backward, 8U of Bst 3.0 DNA polymerase, and 1 µl genomic DNA in a total volume of 25 µl, was incubated at 65°C for 60 min in a T100 Thermal Cycler (Bio-Rad). The amplified products (10 µl) were run on 2% agarose gel and visualized under UV light. For all the tests, a positive control (DNA from a female *Wolbachia*-infected *Ae. albopictus*), a negative control (DNA from females of *Ae. aegypti* Rockefeller strain) and a no template control (nuclease-free water) were used.

### 2.5 Maternal transmission of *Wolbachia* in Las Cruces strain

In August 2017, a colony of *Ae. aegypti* was established from pupae (n=8) collected from a larval habitat in a residential area in Las Cruces. The eclosed adults were confirmed to carry *Wolbachia* by PCR and LAMP assays as described above. Unfortunately, the colony was lost in January 2018. In September 2018, a new colony was initiated from the pupae (n=77) collected from the same location in August 2017. Again, the eclosed adults tested positive for *Wolbachia* The strain was named the LC (Las Cruces, NM) strain. To test whether *Wolbachia* can be transmitted to the progeny maternally, crosses between LC and Rockefeller (Rock) strains were conducted, which included a cross of virgin females (LC) × males (Rock), and a cross of virgin females (Rock) × males (LC). Rockefeller strain of *Ae. aegypti* is known not to carry *Wolbachia* (0% infected out of the 10 individuals screened with LAMP assay). Each cross included 50 females and 30 males of the respective strains. All cages were maintained at 28°C, 72% RH and fed on 20% sucrose till day 3. On day 4, females were blood fed on the forearm of a human volunteer. Eggs were collected at three days post feeding and reared to adults. The progeny adults from the respective crosses were screened for *Wolbachia* by LAMP assay.

## 3 RESULTS

### 3.1 Prevalence of *Wolbachia* in *Ae. aegypti* and *Ae. albopictus* populations in New Mexico

*Aedes aegypti* and *Ae. albopictus* occur in the state of New Mexico (Hahn et al., 2017; Hahn et al., 2016). In 2017, we conducted a survey to map the distribution and characterize the microbiomes of both species in New Mexico. To profile bacteriomes associated with wild *Ae. aegypti* collected from Las Cruces, a bacterial 16S ribosomal RNA gene fragment was amplified by PCR using primers 27F and 519R. The PCR products were cloned and sequenced to identify bacterial taxa, and a total of 80 sequences were obtained from 8 individuals. Unexpectedly, 10 sequences of *Wolbachia* 16S amplicons were identified in six of eight individuals. We then conducted a large survey by screening 148 specimens collected from eight cities across New Mexico, which largely span the distribution of *Ae. aegypti* in the state (Figure 1), by amplifying *Wolbachia gatB* or *ftsZ* (Table 2). Selected positive PCR products were validated by sequencing. Out of 148 specimens tested, 85 were *Wolbachia* positive, the overall prevalence was 57.4% (Tables 2 & 3). *Aedes albopictus* is less common in New Mexico and is only present on the eastern part of the state, and those collected from Clovis (n=12) and Roswell (n=1) were assayed for *Wolbachia* (Table 4). Among the 13 specimens, 11 carried both the *w*AlbA and *w*AlbB strains and one carried wAlbB only. The overall *Wolbachia* prevalence in *Ae. albopictus* in New Mexico was 92.3% (Table 4 & 5). We also amplified DNA fragments of two additional *Wolbachia* genes, encoding PE and DE. The amplicons of *Wolbachia* 16S rDNA, *gatB*, *ftsZ, PE* and *DE* were confirmed by sequencing. In *Ae. aegypti* specimens, only *w*AlbB sequences were detected. Representative sequences were deposited in GenBank (Table S2).

In August 2017, a local *Ae. aegypti* colony was derived from larvae collected in Las Cruces, NM. *Wolbachia* was detected in two of four females and two of four males that eclosed from the pupae. The infection persisted in F1 as well, 14/33 females (42.4%) and 14 of 14 males (100%) of F1 adults examined were positive for *Wolbachia*. Unfortunately, the colony was lost in January 2018 because of a failure of egg hatching at F3 generation. In September 2018, a new colony of *Ae. aegypti* was initiated from the pupae collected from the same location in Las Cruces. The strain, named LC, was *Wolbachia* positive, with a prevalence of 84.6% of the 13 individuals screened.

### 3.2 Prevalence of *Wolbachia* in *Ae. aegypti* and *Ae. albopictus* in Florida

Coon and colleagues (2016) previously reported the presence of *Wolbachia* in the larvae of *Ae. aegypti* collected from Jacksonville, Florida. We therefore examined the specimens of *Ae. aegypti* collected in July 2016 from St. Augustine, Florida, which is approximately 50 miles south of Jacksonville. Among 46 specimens screened, one male and one female were *w*AlbB positive, with a prevalence of 4.3% (Table 3). The *w*AlbA strain was not detected in these specimens. As expected, *Wolbachia* infection occurred at a high prevalence in *Ae. albopictus*. Among the 38 specimens tested, 35 were co-infected with *w*AlbA and *w*AlbB, and one carried only *w*AlbB. The overall *Wolbachia* prevalence in *Ae. albopictus* was 92.1% (Table 5). *Ae. aegypti* and *Ae. albopictus* were sampled at different sites in St. Augustine (Table 1).

### 3.3 Prevalence of *Wolbachia* in *Ae. aegypti* and *Ae. albopictus* in southeastern Texas

In 2018, we screened 98 *Ae. aegypti* and 32 *Ae. albopictus* collected from a neighborhood in Deer Park, Harris County, Texas (Table 1). No *Wolbachia* was detected in *Ae. aegypti*. The overall prevalence of *Wolbachia* infection was 81.3% in *Ae. albopictus*. Among the 26 specimens positive for *Wolbachia*, 15 carried both *w*AlbA and *w*AlbB, two carried *w*AlbA only, and nine carried *w*AlbB only (Table 5).

### 3.4 LAMP assays

The LAMP assay is a sensitive method for detecting low-abundance target DNA in a sample (Notomi et al., 2000). Goncalves Dda et al. (2014) developed a LAMP assay for *Wolbachia* detection in insects (Goncalves Dda, Cassimiro, de Oliveira, Rodrigues, & Moreira, 2014). Recently, Bhadra et al. (2018) reported a specific and sensitive assay that combines LAMP and oligonucleotide strand displacement (OSD) for detecting both species identity and *Wolbachia*. The *Wolbachia* density appears to be low in most infected specimens of *Ae. aegypti*. To corroborate the results from the *Wolbachia* PCR assay, we developed a LAMP assay to detect *Wolbachia*. The LAMP assay was sensitive and detected target *Wolbachia* DNA in infected *Ae. aegypti* directly. As shown in Figure 2, *Wolbachia* positive samples yield a ladder of bands between 200 bp-1kb, and a ∼100 bp band. *Wolbachia* negative samples show an accumulation of oligos around 50 bp. The infected *Ae. albopictus* specimen could be detected at 100 times dilution of template, but not at 500 times dilution, while the infected *Ae. aegypti* specimen could not be detected at 20 times dilution. Figure 2 shows a representative result of LAMP assay on the *Wolbachia* positive and negative specimens from New Mexico, Florida, and Texas.

**Figure 2.**
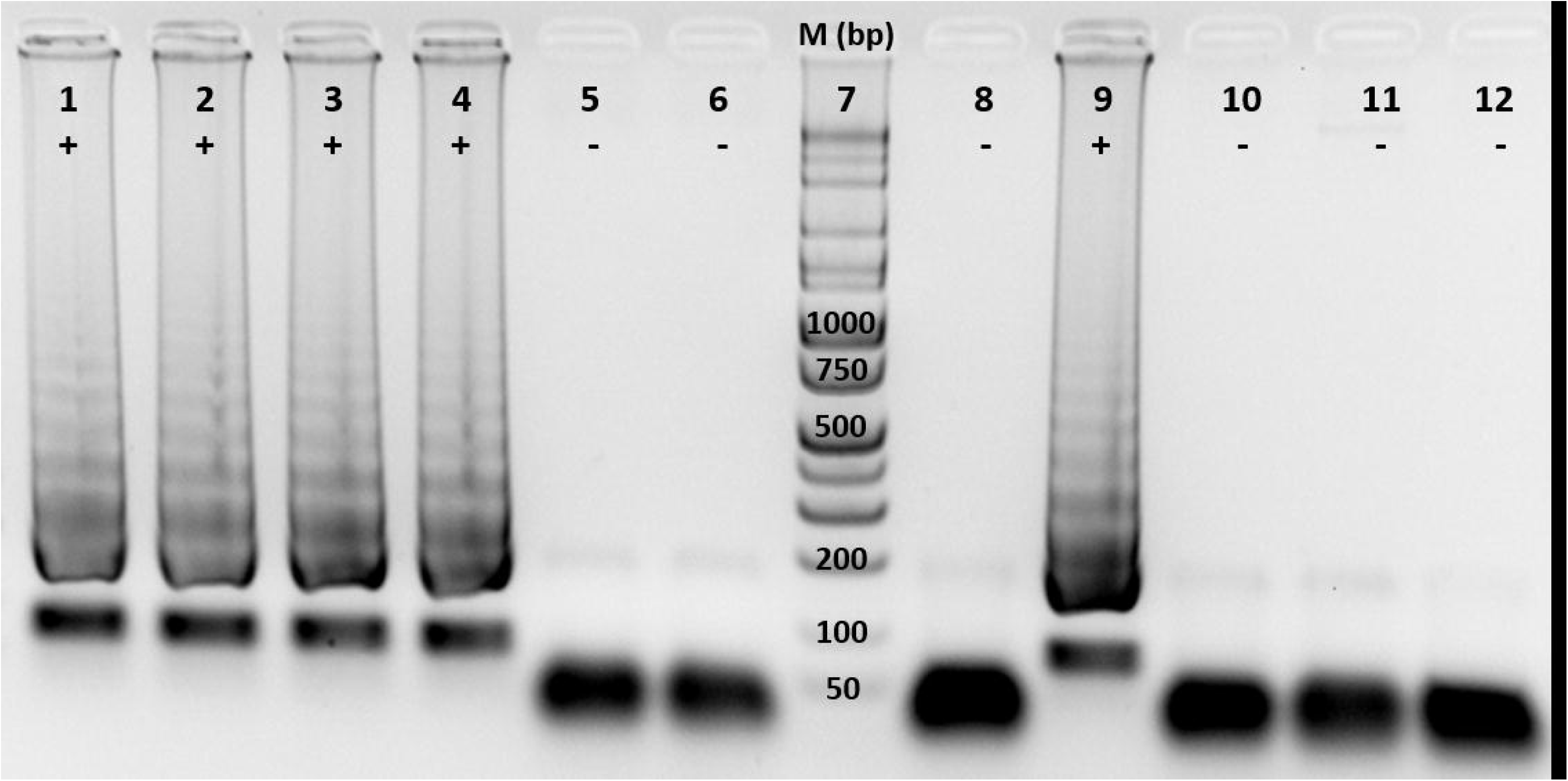
LAMP detection of *Wolbachia* 16S rDNA. 1,2 = *Ae. aegypti* (NM); 3,4 = *Ae. aegypti* (FL); 5,6 = *Ae. aegypti* (TX); 7 = Marker; 8 = *Ae. aegypti* (NM-1, 1:20 dilution); 9 = *Ae. albopictus* (NM, 1:100 dilution); 10 = *Ae. albopictus* (NM, 1:500 dilution); 11 = *Ae. aegypti* Rockefeller; 12 = No template control. +: *Wolbachia* positive; -: *Wolbachia* negative.

### 3.5 Maternal transmission of *Wolbachia* occurs in Las Cruces strain

*Wolbachia* are known to be transmitted vertically from female to the offspring (Werren et al., 2008). To test the occurrence of vertical transmission of *Wolbachia* in LC strain, crosses were set up between the adults of LC strain and Rockefeller strain (Rock). The F2 generation of LC strain was used for the experiment. From the LC females and males used for the crosses, five specimens from each sex were randomly selected and examined by the LAMP assay. They were all *Wolbachia* positive. The crosses between females (LC) × males (Rock) and females (Rock) × males (LC) both yielded viable progenies. The egg hatch rate was significantly higher in F/LC × M/Rock (98/463, 21.2%) than in M/LC × F/Rock (39/475, 8.2%), (Chi square, P<0.01). The progenies were screened for the presence of *Wolbachia* by the LAMP assay. As shown in Figure 3, the progenies of F/LC × M/Rock were *Wolbachia* positive (10/10, 100%), while the progenies of M/LC × F/Rock were *Wolbachia* negative (10/10, 100%). The results clearly demonstrated the maternal transmission of *Wolbachia* in LC *Ae. aegypti*.

**Figure 3.**
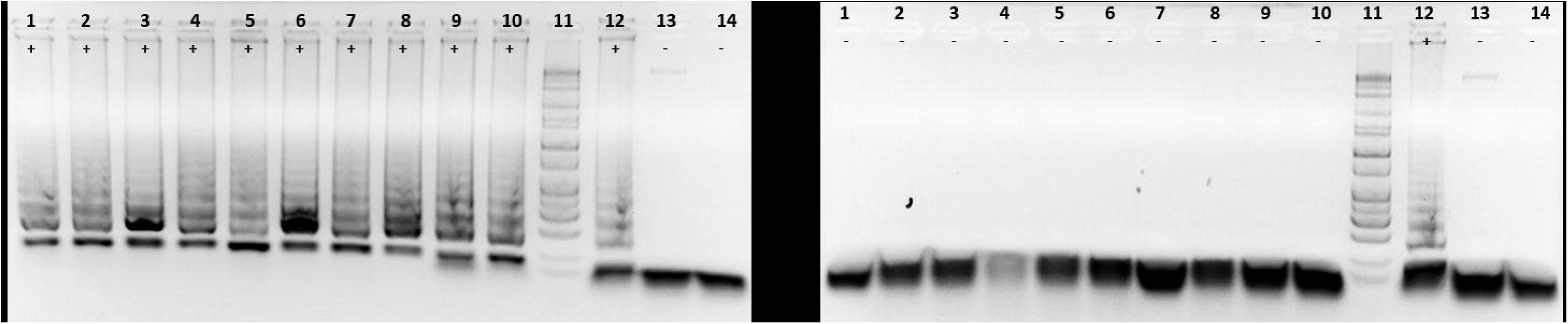
Maternal transmission of *Wolbachia* in LC strain. *Wolbachia* detection in the progeny of the respective crosses. (**A**) 1-10= Progeny of the cross between females (LC) and males (Rock), 11 =Marker, 12 = *Ae. albopictus* (NM, 1:20 dilution), 13= *Ae. aegypti* Rockefeller, 14= No template control. +: *Wolbachia* positive, -: *Wolbachia* negative. **(B)** 1-10= Progeny of the cross between females (Rock) and males (LC), 11 =Marker, 12 = *Ae. albopictus* (NM, 1:20 dilution), 13= *Ae. aegypti* Rockefeller, 14= No template control. +: *Wolbachia* positive, -: *Wolbachia* negative.

## 4 DISCUSSION

*Wolbachia* is commonly associated with wild *Ae. albopictus* around the world. However, *Wolbachia* has not been detected in wild *Ae. aegypti* in previous surveys, until recently. In the study by Coon et al. (2016), two *Wolbachia* 16S rDNA OTUs were detected in a pool of 30 larvae of *Ae. aegypti* that were collected from one of five larval sites in Jacksonville, Florida in June 2014. The *Wolbachia* OTUs were detected in another collection from the same location in July 2014, and both *w*AlbA and *w*AlbB were detected based on the sequence comparison of several *Wolbachia* genes. The prevalence of *Wolbachia* in the *Ae. aegypti* population was not investigated in this study (Coon et al., 2016). Similarly, *Wolbachia* 16S OTUs were detected in two pools of 10 and 25 *Ae. aegypti* respectively, collected in the suburb and urban areas of Thailand (Thongsripong et al., 2017). In a study by Hegde et al. (2018), a small number of *Wolbachia* 16S rDNA reads were found in a few *Ae. aegypti* individuals collected from Houston, Texas, but the results were not validated by PCR using several *Wolbachia* genes. Our survey of Florida mosquitoes was consistent with previous results, detecting a low prevalence (4.3%) of *Wolbachia* in *Ae. aegypti* from St. Augustine, Florida (Table 3). Similarly, no *Wolbachia* was detected in 94 *Ae. aegypti* specimens from Deer Park, Texas.

In contrast, the screening of wild populations of *Ae. aegypti* from eight cities in New Mexico revealed a high prevalence (15.8-100%, average of 57.4%) of *Wolbachia*, a level unprecedented for this species. The infection was validated by sequencing the PCR amplicons from the *ftsZ, gatB, DE and PE* genes (Table S2). These sequences were identical to the sequences of respective genes in the genome of *w*AblB strain (Mavingui et al., 2012), indicating the *Wolbachia* detected in the *Ae. aegypti* samples belongs to the *w*AlbB strain.

A local *Ae. aegypti* strain LC was established from the pupae collected in the fall of 2018. The LC strain was *Wolbachia* positive. LC females were able to pass *Wolbachia* infection to their progeny when crossed with males of *Wolbachia*-free Rockefeller strain (Figure 3), while the infected LC males were did not produce *Wolbachia*- infected offspring when crossed with Rockefeller females, demonstrating maternal transmission. Taken together, the molecular detection of *Wolbachia* in NM *Ae. aegypti* populations and establishment of *Wolbachia* positive colonies in both 2017 and 2018 indicated a persistent transmission of *Wolbachia* with the wild *Ae. aegypti* populations. Furthermore, the maternal transmission of *Wolbachia* to the progeny in the crosses of infected LC and uninfected Rockefeller strains ultimately provided satisfactory evidence that the presence of *Wolbachia* sequences in *Ae. aegypti* reflects authentic infection rather than environmental contamination or *Wolbachia* sequences derived from commensal or parasitic species, such as nematodes, within the mosquitoes.

Recently, Gloria-Soria et al. (2018) reported screening for *Wolbachia* in 2,663 specimens of *Ae. aegypti* from 27 countries, including 60 specimens from Las Cruces, New Mexico, and no *Wolbachia* was detected in these samples. Their screen was conducted on DNA pools of up to 20 individuals. In our survey, we screened individual mosquitoes, and *Wolbachia* density in *Ae. aegypti* was substantially lower than in *Ae. albopictus*. Moreover, most positive specimens were identified by two rounds of PCR. We tested the sensitivity of our PCR assay using a DNA pool comprising 19 *Wolbachia* negative individuals and one positive individual for both *Ae. aegypti* and *Ae. albopictus*. *Wolbachia* could be detected in the pool containing DNA from the single positive *Ae. albopictus* specimen, but not in the pool containing DNA from the single positive *Ae. aegypti* specimen (data not shown). *Wolbachia* titer shows striking variation in infected individuals of *Ae. albopictus* (Ahantarig, Trinachartvanit, & Kittayapong, 2008; Calvitti, Marini, Desiderio, Puggioli, & Moretti, 2015). For example, in wild-caught *Ae. albopictus* in North and Central Italy, 30.8-50.0% of infected male *Ae. albopictus* had a low titer of *w*AlbA, which was not detectable by a standard PCR, but detectable by a quantitative PCR assay, while in the remaining males having higher *Wolbachia* densities it was detectable by a standard PCR (Calvitti et al., 2015). Similarly, *Wolbachia* load also varies substantially in *Ae. aegypti* into which the bacterium has been artificially transinfected (Ant, Herd, Geoghegan, Hoffmann, & Sinkins, 2018). The *Wolbachia* load was quite low in the *Ae. aegypti* samples in our study. Therefore, we hypothesize that assay sensitivity explains the different results between our study and that of Gloria-Soria et al. (2018).

The conspicuous variation in the prevalence of *Wolbachia* infection in different populations of *Ae. aegypti* among the three states and eight cities within New Mexico raises several compelling questions. Chief among them are ‘what factors contribute to the low density of *Wolbachia* in *Ae. aegypti* relative to *Ae. albopictus?’*, and ‘what makes the New Mexico populations more susceptible/hospitable to *Wolbachia* than other populations?’

*Wolbachia* density variation is common in natural populations of different insect hosts (Ahantarig et al., 2008; Unckless, Boelio, Herren, & Jaenike, 2009), however, it is not clear whether the variation is driven by genetic or environmental variation or both. A recent study revealed an amplification of a genome region that harbors a cluster of eight genes, called Octomom, which is responsible for the over-replication and virulence of *w*MelPop in *Drosophila melanogaster.* The copy number of Octomom correlates with *Wolbachia* titers (Chrostek & Teixeira, 2015). Environmental factors such as temperature may play a role as well. A recent study in *D. melanogaster* demonstrated that *Wolbachia* abundance was higher when host flies developed at lower temperature (13°C and 23°C compared to 31°C) (Moghadam et al., 2018). Additionally, mosquito genetic background may affect *Wolbachia* prevalence. It has been shown that *Ae. aegypti* populations from Las Cruces, New Mexico, Houston, Texas, and four locations of Florida are genetically distinct (Pless et al., 2017). Further studies are needed to tease out the roles of these potential drivers of *Wolbachia* presence and abundance in *Ae. aegypti*.

Another critical question raised by our study is how *Ae. aegypti* acquires a strain of *Wolbachia* associated with *Ae. albopictus* and how this strain impacts *Ae. aegypti* life history and pathogen susceptibility. *Ae. aegypti* can be artificially transinfected with different *Wolbachia* strains, and the artificial infection can be introduced into natural populations and spread in nature (Frentiu et al., 2014; Hoffmann et al., 2011; Schmidt et al., 2017). The *w*AlbB has been successfully introduced into *Ae. aegypti* to form a line with inherited infection (Xi, Khoo, & Dobson, 2005). Interestingly, the Toll and IMD pathways favor establishment and maintenance of *w*AlbB infection in the line; the knockdown of Toll and IMD by RNA interference reduces the *w*AlbB load, while the transgenic activation of Toll and IMD increases the load (Pan et al., 2018). It appears that transinfected *Wolbachia* can exploit host immunity for a symbiotic association. Our survey revealed the prevalence of *w*AlbB in *Ae. aegypti* natural populations in New Mexico, and an infected colony has been established from wild-collected mosquitoes. This provides an opportunity to study the natural *Ae. aegypti-Wolbachia* association and its impact on various mosquito life traits, such as reproductive manipulation, as well as interference with viral transmission.

## ACKNOWLEDGMENTS

We thank the participants of SouthWest *Aedes* Research and Mapping project (SWARM); Jennifer Corrado at Anastasia Mosquito Control District in St. Augustine, FL; and Elaine Chu at Harrison County Public Health, Mosquito and Vector Control Division in Houston, TX for their contributions to the mosquito sampling. This work was supported by New Mexico Department of Health contract to Kathryn A. Hanley, Michaela Buenemann, Immo A. Hansen and Jiannong Xu. Concepcion Sanchez was an undergraduate research scholar supported by the Institutional Development Award (IDeA) from the National Institute of General Medical Sciences of the National Institutes of Health under grant number P20GM103451. The funder had no roles in the study design, data collection and analysis, or decision to publish. The content is solely the responsibility of the authors and does not necessarily represent the official views of the National Institutes of Health and the Department of Health of New Mexico.

## AUTHORS’ CONTRIBUTIONS

JX conceived the study design. KJLM, DD, KH and MD collected mosquitoes. AK, WY, JJ, CS AKK conducted assays and data analysis. JX, AK, MB and AKK drafted the manuscript, KAH, DD, MD, MB, IAH and RX critically reviewed the manuscript. All authors read and approved the manuscript.

## DATA ACCESSEBILITY

DNA sequences were deposited in GenBank under accession number MH732668-MH732670, MH734116-MH734121.

## Additional files

**Table S1. Primer sets used in the study**

**Table S2. GenBank accession numbers of Wolbachia sequences**

