## Supplementary material for "*Wolbachia pipientis* occurs in *Aedes aegypti* populations in New Mexico and Florida, USA": Table 5

**Table 5. *Wolbachia* strain distribution in *Aedes albopictus* in Texas, Florida, and New Mexico**

| Specimens (n) | *Wolbachia* strain | | | |
| --- | --- | --- | --- | --- |
|  | A & B (%) | A (%) | B (%) | Total No. (%) |
| Male (19), TX | 6 (31.6) | 1 (5.3) | 9(47.4) | 16 (84.2) |
| Female (13), TX | 9 (69.2) | 1 (7.7) | 0 | 10 (76.9) |
| Male (20), FL | 20 (100) | 0 | 0 | 20 (100) |
| Female (18), FL | 14 (77.8) | 0 | 1 (5.6) | 15 (83.3) |
| Male (2), NM | 0 | 0 | 1(50.0) | 1(50.0) |
| Female (11), NM | 11(100) | 0 | 0 | 11(100) |
