## Supplementary material for "*Wolbachia pipientis* occurs in *Aedes aegypti* populations in New Mexico and Florida, USA": Table 1

**Table 1. Mosquito collections in Florida, New Mexico and Texas**

| Species | Location | Coordinates | Collection time |
| --- | --- | --- | --- |
| *Ae. aegypti* | St. Augustine, FL | 29.895, -81.313 | July, 2016 |
| *Ae. albopictus* | St. Augustine, FL | 29.890, -81.332 | July, 2016 |
| *Ae. aegypti* | Deer Park, TX | 29.693, -95.115 | May, 2018 |
| *Ae. albopictus* | Deer Park, TX | 29.693, -95.115 | May, 2018 |
| *Ae. aegypti* | 8 cities, NM | see Table 2 | May-Nov, 2017 |
| *Ae. albopictus* | 2 cities, NM | see Table 4 | May-Nov, 2017 |
