## Supplementary material for "*Wolbachia pipientis* occurs in *Aedes aegypti* populations in New Mexico and Florida, USA": Table 2

**Table 2. *Wolbachia* infection in *Aedes aegypti* populations in New Mexico from May-November 2017**

| City (n) | Infection rate (%) | Coordinates of collection sites |
| --- | --- | --- |
| Alamogordo (19) | 3 (15.8) | 32.861, -105.979; 32.918, -105.936 |
| Carlsbad (31) | 6 (19.4) | 32.356, -104.248; 32.440, -104.240; 32.427, -104.223 |
| Deming (29) | 26 (89.7) | 32.251, -107.763; 32.245, -107.761; 32.262, -107.745 |
| Las Cruces (30) | 24 (80.0) | 32.296, -106.732; 32.357, -106.769; 32.396, -106.816 |
| Lovington (9) | 7 (77.8) | 21.491, -103.364 |
| Sunland Park (2) | 2 (100) | 31.816, -106.603 |
| Roswell (26) | 16 (61.5) | 33.378, -104.513; 33.416, -104.529 |
| Truth or Consequences (2) | 1 (50.0) | 33.120, -107.272; 33.203, -107.228 |
| Total (148) | 85 (57.4) |  |
