## Supplementary material for "*Wolbachia pipientis* occurs in *Aedes aegypti* populations in New Mexico and Florida, USA": Table 3

**Table 3. *Wolbachia* infection in *Aedes aegypti* in New Mexico and Florida**

| Specimens (n) | *Wolbachia* strain | | | |
| --- | --- | --- | --- | --- |
|  | A & B (%) | A (%) | B (%) | Total No. (%) |
| Male (51), NM | 0 | 0 | 28(54.9) | 28(54.9) |
| Female (97), NM | 0 | 0 | 57(58.8) | 57(58.8) |
| Male (18), FL | 0 | 0 | 1(5.5) | 1(5.5) |
| Female (28), FL | 0 | 0 | 1 (3.6) | 1(3.6) |
