## Supplementary material for "*Wolbachia pipientis* occurs in *Aedes aegypti* populations in New Mexico and Florida, USA": Table 4

**Table 4. *Wolbachia* infection in *Ae. albopictus* populations in New Mexico from May-November 2017**

| City (n) | Infection rate (%) | Coordinates of collection sites |
| --- | --- | --- |
| Clovis (12) | 11 (91.7) | 34.406, -103.192; 34.424, -103.182; 34.414, -103.196; 34.399, -103.200; 34.404, -103.201 |
| Roswell (1) | 1 (100) | 33.389, -104.530 |
| Total (13) | 12 (92.3) |  |
