## Supplementary material for "*Wolbachia pipientis* occurs in *Aedes aegypti* populations in New Mexico and Florida, USA": Table S1

| **Table S1. Primer sets used in the study** | | | |
| --- | --- | --- | --- |
| *Species identification PCR* | | Annealing Tm | Reference |
| 18SFHIN | GTAAGCTTCCTTTGTACACACCGCCCGT | 55 | Higa et al. (2010) |
| aeg.r1 | TAACGGACACCGTTCTAGGCCCT |  |  |
| alb.r1 | GTACTAGGCTCACTGCCACTGA |  |  |
| *Bacterial 16S rRNA gene* | |  |  |
| 27F | GAGTTTGATCNTGGCTCAG | 50 | Wang et al. (2011) |
| 519R | GTNTTACNGCGGCKGCTG |  |  |
| *gatB* |  |  | Baldo et al. (2006) |
| wAlbB_gatB_F | TAAGAATCGCAAGAATTCAC | 50 |  |
| wAlbB_gatB_R | TGGYAAYTCRGGYAAAGATGA |  |  |
| wAlbA_gatB_F | TTTAGAGCAAGATGCAGGRAAGAGCG | 50 |  |
| wAlbA_gatB_R | TGGYAAYTCRGGYAAAGATGA |  |  |
| *fitsZ* |  |  | Baldo et al. (2006) |
| wAlbB_ftsZ_F | AAAGATAGCCATATGCTCTTT | 50 |  |
| wAlbB_ftsZ_R | CATTGCTTTACCCATCTCA |  |  |
| wAlbA_ftsZ_F | AAAGATAGTCATATGCTTTTC | 50 |  |
| wAlbA_ftsZ_R | CATCGCTTTGCCCATCTCG |  |  |
| *Phosphoesterase* | |  | This study |
| wAlbB_PE_F | CGCAGCTCAATTAACAATACC | 55 |  |
| wAlbB_PE_R | CAATCAGCGTACCAAGCTTCT |  |  |
| wAlbA_PE_F | ACTCAATTAACAATGGTGATCAT | 55 |  |
| wAlbA_PE_R | AGCTCTCTTACAACTCTCATGG |  |  |
| *Diaminopimelate epimerase (EC 5.1.1.7)* | |  | This study |
| wAlbB_DE_F | TGTGAATGCGATCCCCTTTATC | 55 |  |
| wAlbB_DE_R | TATCGCCAGTCATAAATATATTTCC | |  |
| wAlbA_DE_F | ACTGAGTGTGACACTCTTCACT | 55 |  |
| wAlbA_DE_R | CTGTTGGGCAGTCAGATATCC |  |  |
| LAMP, 16S ribosomal RNA gene | | N/A | This study |
| F3 | CTGGAACTGAGATACGGTC |  |  |
| B3 | TTACGCCCAATAATTCCGA |  |  |
| FIP | TCTTCACTCATGCGGCATGGCAGTGGGGAATATTGGACAA | | |
| BIP | AGGAAGATAATGACGGTACTCACAGATAACGCTAGCCCTCTCC | | |
| LF | CTGGATCAGGCTTTCGCCC |  |  |
| LB | AGTCCTGGCTAACTCCGTG |  |  |
