## Supplementary material for "*Wolbachia pipientis* occurs in *Aedes aegypti* populations in New Mexico and Florida, USA": Table S2

**Table S2. GenBank accession numbers of *Wolbachia* sequences**

| Sequence ID | Host mosquito | *Wolbachia* gene | *Wolbachia* strain | GenBank accession |
| --- | --- | --- | --- | --- |
| S77 | *Ae. aegypti* | 16S rRNA | N/A | MH732668 |
| S56 | *Ae. aegypti* | 16S rRNA | N/A | MH732669 |
| S66 | *Ae. aegypti* | 16S rRNA | N/A | MH732670 |
| LC40 | *Ae. aegypti* | ftsZ | *w*AlbB | MH734116 |
| D4a | *Ae. aegypti* | gatB | *w*AlbB | MH734120 |
| LC11 | *Ae. albopictus* | diaminopimelate epimerase (DE) | *w*AlbA | MH734119 |
| FL139 | *Ae. aegypti* | diaminopimelate epimerase (DE) | *w*AlbB | MH734118 |
| TX120 | *Ae. albopictus* | phosphoesterase (PE) | *w*AlbA | MH734121 |
| FL8 | *Ae. aegypti* | phosphoesterase (PE) | *w*AlbB | MH734117 |
